# Patient-Specific Heart Rate Modulates Developmental Electrophysiology in Transcriptomic-Guided In Silico Models of Pediatric Human Atrial Cardiomyocytes

**DOI:** 10.64898/2026.07.21.739866

**Authors:** Gabriella M. Ellks, Mario J. Mendez, Devon Guerrelli, Jacob A. Miller, Manan Desai, Yves d’Udekem, Nikki Gillum Posnack, Seth H. Weinberg

**Affiliations:** Department of Biomedical Engineering, The Ohio State University, Columbus, OH; Children’s National Heart and Lung Institute, Children’s National Hospital, Washington, DC; Sheikh Zayed Institute for Pediatric Surgical Innovation, Children’s National Hospital, Washington, DC; Department of Biomedical Engineering, The George Washington University, Washington, DC; Division of Cardiovascular Surgery, Children’s National Hospital, Washington, DC; Department of Pharmacology and Physiology, The George Washington University, Washington, DC; Department of Pediatrics, The George Washington University, Washington, DC; Davis Heart and Lung Research Institute, The Ohio State University, Columbus, OH

**Author notes:** Corresponding author: Seth H. Weinberg.

**Keywords:** Computational modeling, Population modeling, Transcriptomics, Electrophysiology, Pediatrics

## Abstract

Cardiac electrophysiology adapts throughout pediatric development, driven by factors including age-associated ion channel expression changes and decreasing heart rate. Our prior transcriptomic-guided simulations of human atrial cardiomyocytes predicted developmental-associated changes in electrophysiology biomarkers at a fixed pacing rate, leaving the contribution of age- and patient-specific heart rate unresolved. In this study, we incorporated intrinsic heart rate into gene expression-guided computational models to predict the interaction between developmental maturation and pacing rate to shape atrial electrophysiology. Virtual patient-specific populations of atrial cardiomyocytes were generated from the right atrial cardiomyocyte gene expression data from 117 patients, spanning neonates to young adults. We simulated each population at pacing rates corresponding to each patient’s intrinsic ECG-based heart rate and at fixed rates corresponding to the patient cohort minimum, median, and maximum. Action potential and calcium transient biomarkers were quantified, and partial least squares regression assessed key biomarker dependencies. For intrinsic-rate pacing conditions, action potential duration at 50% and 90% repolarization increased with age, whereas early repolarization shortened; maximum upstroke velocity increased, resting membrane potential became more negative, and alternans prevalence decreased. Developmental differences persisted during fixed-rate pacing conditions, indicating that differences were not explained solely by the faster heart rates of younger patients. Notably, intrinsic-rate simulations exhibited stronger age associations for upstroke velocity and alternans than fixed-rate simulations. Sensitivity analyses indicated that electrophysiological phenotypes arose from interactions among ionic conductances, calcium handling, age, and heart rate. Collectively, we find that pediatric atrial electrophysiology reflects both intrinsic developmental remodeling and rate-dependent modulation.

## Introduction

Congenital heart disease (CHD) remains the leading cause of mortality from birth defects and continues to impose a substantial disease burden worldwide, despite major advances in cardiovascular medicine and surgery (1). Although advances in pediatric cardiac surgery and care have improved survival, younger age at the time of surgery remains associated with increased risk of postoperative complications, including arrhythmias, myocardial injury, and impaired mechanical function (2–5). These age-dependent vulnerabilities are likely due to the structural, metabolic, and electrophysiological immaturity of the developing myocardium (6). A more complete understanding of postnatal cardiac maturation is therefore essential for improving age-appropriate clinical decision-making, including surgical timing, myocardial protection strategies, postoperative management, and pharmacological therapy.

During postnatal development, cardiomyocytes undergo coordinated remodeling that supports the transition from the intrauterine to extrauterine environment. These changes include a shift from proliferative to hypertrophic growth, maturation of sarcomeric organization, remodeling of excitation-contraction coupling, development of intercellular electrical and mechanical connections, and age-dependent changes in ion channel expression and function (7–18). In humans, several aspects of cardiomyocyte maturation continue beyond infancy, with structural and electrophysiological properties evolving over the first years of life and, in some cases, into childhood (7,13). However, much of the current understanding of postnatal cardiac maturation is derived from animal models, due in part to the limited availability of pediatric human cardiac tissue (19,20). Existing human studies have provided important insight into developmental changes in contractile kinetics, calcium handling, and action potential morphology, but many have been constrained by small sample sizes, limited age stratification, or the inability to integrate molecular and functional measurements within the same framework (10,12,14–18).

Developmental remodeling of cardiac electrophysiology is particularly relevant in pediatric patients, because immature cardiomyocytes may respond differently to physiological stress, surgical injury, autonomic stimulation, pacing rate, and ion channel-targeting medications. Prior studies have reported age-dependent differences in human atrial action potential morphology, calcium current properties, and transient outward potassium current density (14–18). In parallel, heart rate decreases substantially from infancy to adulthood, introducing an additional variable that may influence developmental electrophysiology (21). As action potential duration, calcium cycling, repolarization reserve, and alternans exhibit well-established rate-dependence (22,23), age-related differences in atrial electrophysiology may reflect both the intrinsic molecular maturation and physiological pacing environment in which immature cardiomyocytes operate.

Recent advances in transcriptomics and computational modeling provide an opportunity to connect molecular remodeling with predicted electrophysiological function. In a recent study, we collected right atrial tissue from pediatric and young adult patients with acyanotic CHD and demonstrated age-dependent adaptations in cardiac gene expression, including genes associated with ion channels, calcium handling, structural organization, metabolism, and cell cycle regulation (24). By integrating these transcriptomic data with a human atrial cardiomyocyte computational model, we predicted developmental differences in action potential and calcium transient morphology and identified ionic current changes associated with postnatal electrophysiological maturation (24–27). However, to avoid introducing frequency-dependent effects, simulations in our prior study were performed at a fixed pacing rate of 60 beats/min.

As a result, the extent to which patient-specific intrinsic heart rate contributes to simulated developmental electrophysiology remains unresolved. This distinction is important as developmental changes in gene expression and developmental changes in heart rate occur in parallel. Thus, comparing the responses from simulations performed at the intrinsic (patient-specific) pacing rate with fixed pacing rate simulations provides a means to distinguish developmental changes that persists across pacing conditions (i.e., are rate-independent) from electrophysiological differences that are modulated by heart rate.

In the present study, we addressed this limitation by incorporating patient-specific intrinsic heart rate into transcriptomic-guided simulations of pediatric human atrial cardiomyocytes. Using previously collected right atrial gene expression data from pediatric and young adult patients with acyanotic CHD (24), we generate patient-specific virtual populations by scaling ionic conductances according to normalized gene expression levels. We then simulate action potentials and calcium transients at each patient’s intrinsic heart rate and at fixed pacing rates spanning the observed cohort range. The main objective of this study was to determine how developmental age and intrinsic heart rate interact to shape simulated atrial electrophysiology across pediatric maturation. We hypothesized that age-dependent electrophysiological differences would persist across pacing conditions, reflecting underlying developmental remodeling, but that intrinsic heart rate would modulate the magnitude of these differences and contribute to variation in repolarization, calcium handling, and alternans susceptibility.

## Methods

### Human Tissue Collection and Gene Expression Profiling

This study used previously collected right atrial tissue and gene expression data from pediatric and young adult patients undergoing cardiac surgery at Children’s National Hospital (24). Tissue collection was performed under a Children’s National Hospital Institutional Review Board-approved protocol (24), and right atrial tissue was collected as surgical waste from patients with acyanotic congenital heart disease and oxygen saturation greater than 94%. The original cohort included 117 patients ranging from 5 days to 32 years of age and was stratified into five developmental age groups: neonate, infant, toddler/preschool, school age, and adolescent/young adult (Supplemental Table 1). Clinical data collected and used in this study for each patient included age, heart rate, and sex.

Gene expression profiling was performed in the original study using RNA isolated from right atrial tissue and microarray-based transcriptomic analysis. Differentially expressed genes were previously identified across developmental age groups using one-way ANOVA with thresholds of P ≤ 0.05, absolute fold change ≥ 1.25, and false discovery rate ≤ 0.1 (24). In the present study, normalized gene expression values from this prior dataset were used to generate patient-specific ionic conductance scaling factors for computational simulations.

### Computational Modeling Framework

#### Patient-Specific Virtual Population Generation

Patient-specific simulations were generated using normalized gene expression values from the prior transcriptomic dataset. As in the original study (24), gene expression values corresponding to ionic currents and calcium-handling components were normalized relative to the adolescent/young adult cohort and used as conductance scaling factors in a human atrial cardiomyocyte model. The original adult control population was generated by varying 18 ionic currents and exchangers between 0.25- and 4-fold of baseline using Latin hypercube sampling. Simulated cells were retained if their action potential (AP) and calcium transient (CaT) biomarkers were within experimentally reported ranges for adult right atrial tissue, resulting in a fitted control adult population of 155 atrial cell models.

For the present study, patient-specific virtual population were generated by multiplying each patient’s normalized gene expression for ionic current by each member of the 155-cell control population. This produced 155 simulated atrial cells per patient, preserving both patient-level gene expression differences and intercellular variability from the fitted adult control population.

#### Heart-Rate-Specific Simulations

Atrial cardiomyocyte electrophysiology was simulated using the Morotti-Grandi human atrial cardiomyocyte model implemented in MATLAB (25,26). Our primary analysis focused on performing patient-specific populations, with each simulation using a pacing basic cycle length (BCL) corresponding with the specific patient’s average heart rate. For outpatients, heart rate values were obtained from their most recent pre-surgical cardiology clinic visit, vitals the morning of surgery, and/or heart rate on arrival to the operating room. For inpatients, heart rate values were obtained from continuous monitoring in the hospital, vitals the morning of surgery, and/or heart rate on arrival to the operating room. For comparison, three additional simulation sets were performed using fixed heart rates corresponding to the minimum, median, and maximum average heart rates observed in the patient cohort: 64, 119, and 171 beats/min, respectively. These fixed-rate simulations were generated using the same patient-specific virtual populations, with only the pacing BCL altered between conditions. For each simulation and pacing rate, the stimulus threshold was determined for each patient-specific cell model using a bisection approach, with complete capture defined as successful depolarization of the final 10 paced beats. The stimulus amplitude was then set to 1.5 times the calculated capture threshold.

#### Biomarker Measurement

AP biomarkers were calculated from the simulated membrane voltage and included APD30, APD50, APD90 (action potential duration at 30%, 50%, and 90% repolarization), diastolic interval, peak voltage, action potential amplitude (APA), resting membrane potential (RMP), action potential triangulation (Tri90-50), plateau potential at 20% of APD90 (PLT20), and maximum upstroke velocity (dV/dt_max_). Calcium transient biomarkers were calculated from the simulated intracellular calcium transient and included CaT amplitude, peak calcium, minimum calcium before the peak, and the timing of peak and minimum calcium. Each cell model was paced to steady state before for a minimum of 100 beats and maximum of 1000 beats, with steady state assessed every 25 beats and defined as less than 0.1% change in both APD50 and APD90 over the final 10 beats, or until the maximum number of paced beats was reached.

For each simulated cell, APD30, APD50, APD90, AP amplitude, dV/dt_max_, plateau potential, and triangulation were calculated as the mean value across the analyzed beats. RMP was calculated as the maximum value from the resting membrane potential output, and calcium transient amplitude was calculated as the maximum calcium transient amplitude across analyzed beats. After individual simulation level biomarker extraction, the patient-specific population biomarkers were defined as the median of the 155 simulated cells.

#### Alternans Analysis

Alternans were assessed using the final two simulated beats for each patient-specific cell simulation. APD50 and APD90 alternans magnitude was calculated as the difference between the long and short values. The presence of APD50 and APD90 alternans were classified by a beat-to-beat difference of at least 5 ms. CaT amplitude alternans was calculated as 1-(CaAmp_small_/CaAmp_large_), and was classified as present when this normalized value was at least 0.3. For each patient, the number and percentage of simulated cells displaying APD50 alternans, APD90 alternans, or calcium transient amplitude alternans were calculated across the 155-cell virtual population.

#### Partial Least Squares Regression Analysis

Partial least squares regression was used to evaluate relationships between patient-level inputs and simulated electrophysiological outcomes. The predictor matrix included patient age, patient average heart rate, and normalized gene expression-derived conductance scaling factors. The response matrix consisted of patient-level summarized biomarkers, including APD30, APD50, APD90, action potential amplitude, maximum upstroke velocity, resting membrane potential, plateau potential, triangulation, CaT amplitude, and the patient-level percentages of simulated cells exhibiting APD50 alternans, APD90 alternans, and CaT amplitude alternans.

#### Statistical Analysis for Age-Dependent Biomarker Comparisons

For age-group comparisons, patients were assigned to one of five developmental groups: neonate, infant, toddler/preschool, school age, or adolescent/young adult. Patient-level biomarker values were compared across age groups using either one-way ANOVA or Kruskal–Wallis testing, depending on distributional assumptions. For each biomarker, normality was assessed within each age group using the Lilliefors test, and equality of variance across groups was assessed using Levene’s absolute test. If all age groups were normally distributed and variances were equal, one-way ANOVA was performed. If either assumption was not met, Kruskal–Wallis testing was used. Post hoc multiple comparisons were performed using Dunn–Šidák correction. Statistical significance was defined as P < 0.05. In the plotted age-group comparisons, each younger age group was compared with the adolescent/young adult reference group.

Age-dependent trends were also evaluated using linear regression, using the log-transformed age (in days). For each biomarker, a linear model was fit between log-transformed age and the patient-level biomarker value, with the coefficient of determination, R², and slope-associated p-value reported.

## Results

### Study Cohort and Age-Dependent Heart Rate Trends

The study cohort included pediatric patients spanning five developmental stages: neonate, infant, toddler/preschool, school age, and adolescent/young adult. Patient intrinsic heart rate demonstrated the expected inverse relationship with age (Figure 1), with the highest rates observed in neonates and infants and progressively slowing through adolescence. However, substantial inter-individual variability in intrinsic heart rate was present within each age group.

**Figure 1.**
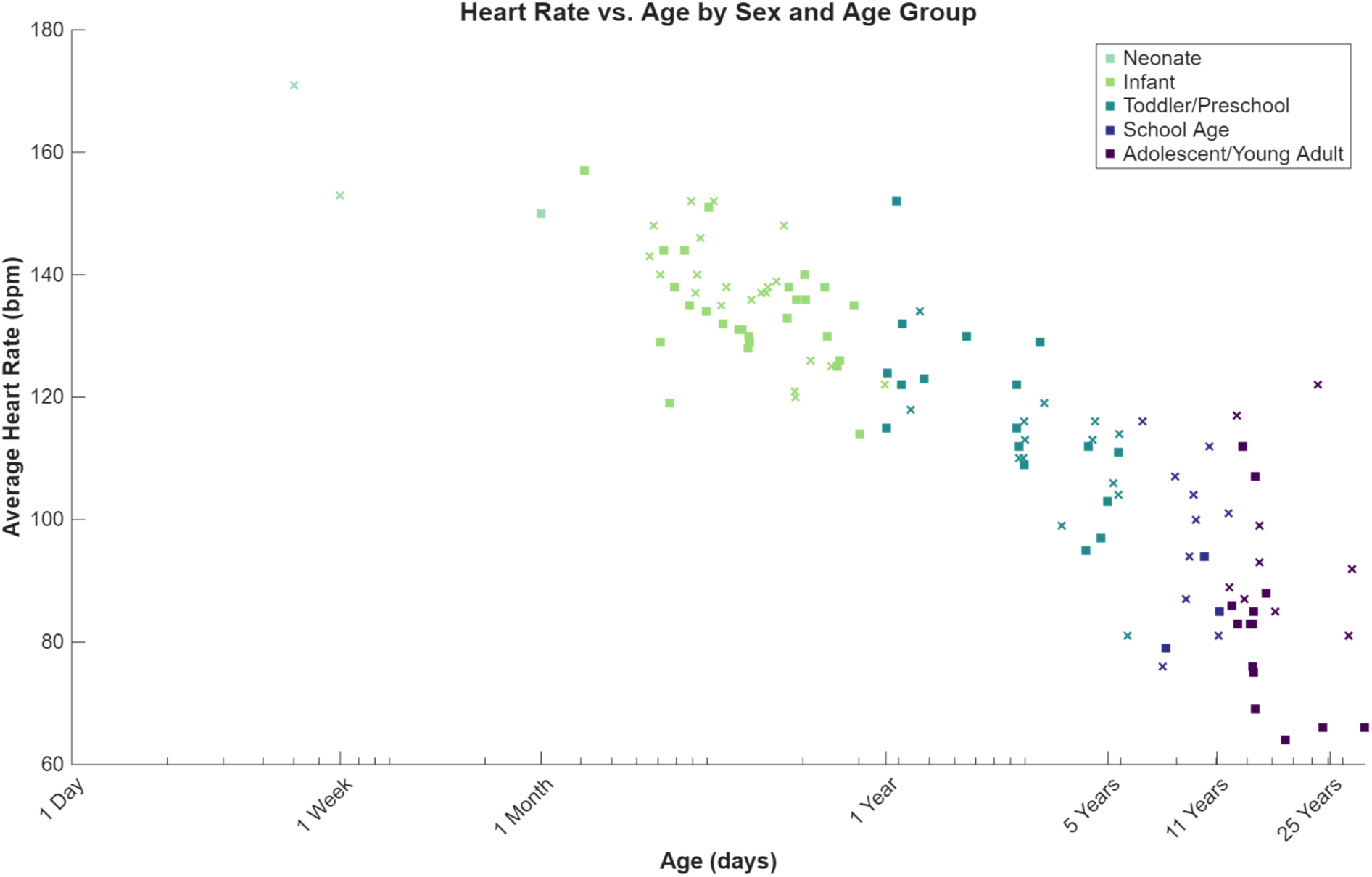
Scatter plot of the average heart rates (beats per minute) versus age (days) for individual patients, stratified by pediatric age group and sex. Colors represent age groups (Neonate, Infant, Toddler/Preschool, School Age, and Adolescent/Young Adult), and marker indicates sex (males denoted by “□”, females by “×”).

### Age-Dependent Differences in Simulated Electrophysiology

#### Representative Action Potentials and Calcium Transients

As described above, patient-specific populations of cardiomyocytes were generated by scaling ionic conductances based on gene expression. Simulations revealed distinct, age-dependent differences in both action potential morphology and CaT dynamics (Figure 2). Younger age groups exhibited shorter repolarization phases and altered calcium handling compared with adolescent/young adult models, demonstrating that developmental stage and heart rate directly influence electrophysiological phenotype.

**Figure 2.**
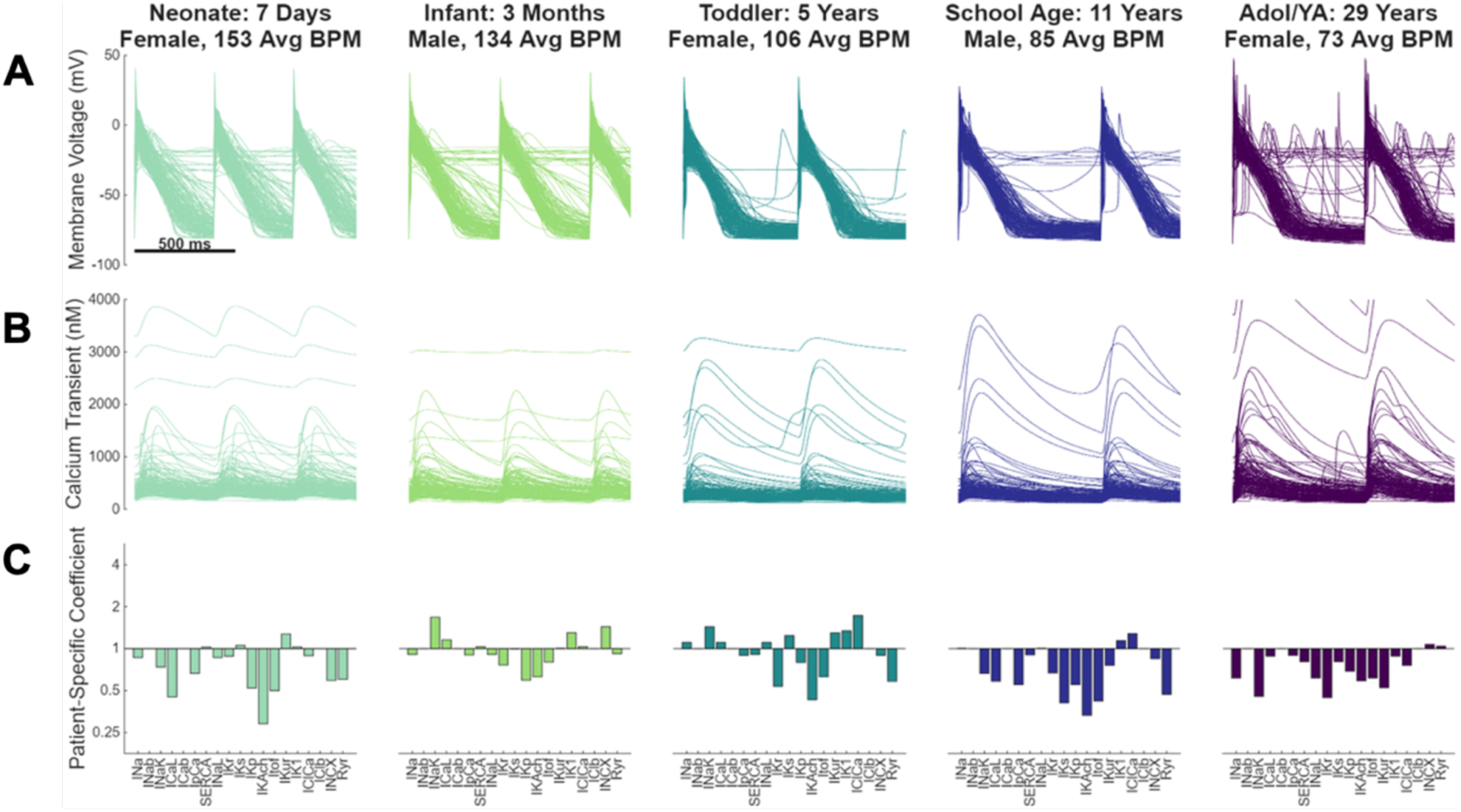
Age-dependent simulated action potentials, calcium transients, and patient-specific conductance coefficients. A: Simulated action potentials from representative patient-specific populations of cells for each of the five age groups (Neonate, Infant, Toddler/Preschool, School Age, and Adolescent/Young Adult) paced at the intrinsic heart rate of the patient (500 ms scale bar). B: Simulated calcium transients from representative patient-specific populations of cells for each age group paced at the patient’s intrinsic heart rate based on same scale. C: Simulations are based on patient-specific gene expression, normalized to the median value of the young adult cohort, representing patient-specific current conductance coefficients.

#### Developmental Trends in Electrophysiological Biomarkers

We next quantified electrophysiological biomarkers for each age group, with each patient stimulated at a pacing rate matching their intrinsic heart rate (Figure 3). We found that APD50 and APD90 both increased progressively with age, whereas early repolarization (APD30) decreased with age. The maximum depolarization rate (dV/dt_max_) increased with age, consistent with enhanced excitability in more mature cardiomyocytes. Action potential triangulation (Tri90-50) also increased across development, reflecting a more gradual repolarization profile in older groups, while resting membrane potential (RMP) became more hyperpolarized. Further, the prevalence of alternans in both action potential and calcium transient prevalence decreased with age. Statistical comparisons identified significant differences between younger age groups and the adolescent/young adult reference group, particularly in infants and toddlers, for several key biomarkers (i.e., APD30, APD90, APA, dV/dt_max_, Tri90-50). Regression analyses based on the medians for each individual patient demonstrated log-linear relationships between age and multiple electrophysiological biomarkers, with the strongest associations observed for APD90, dV/dt_max_, and alternans prevalence.

**Figure 3.**
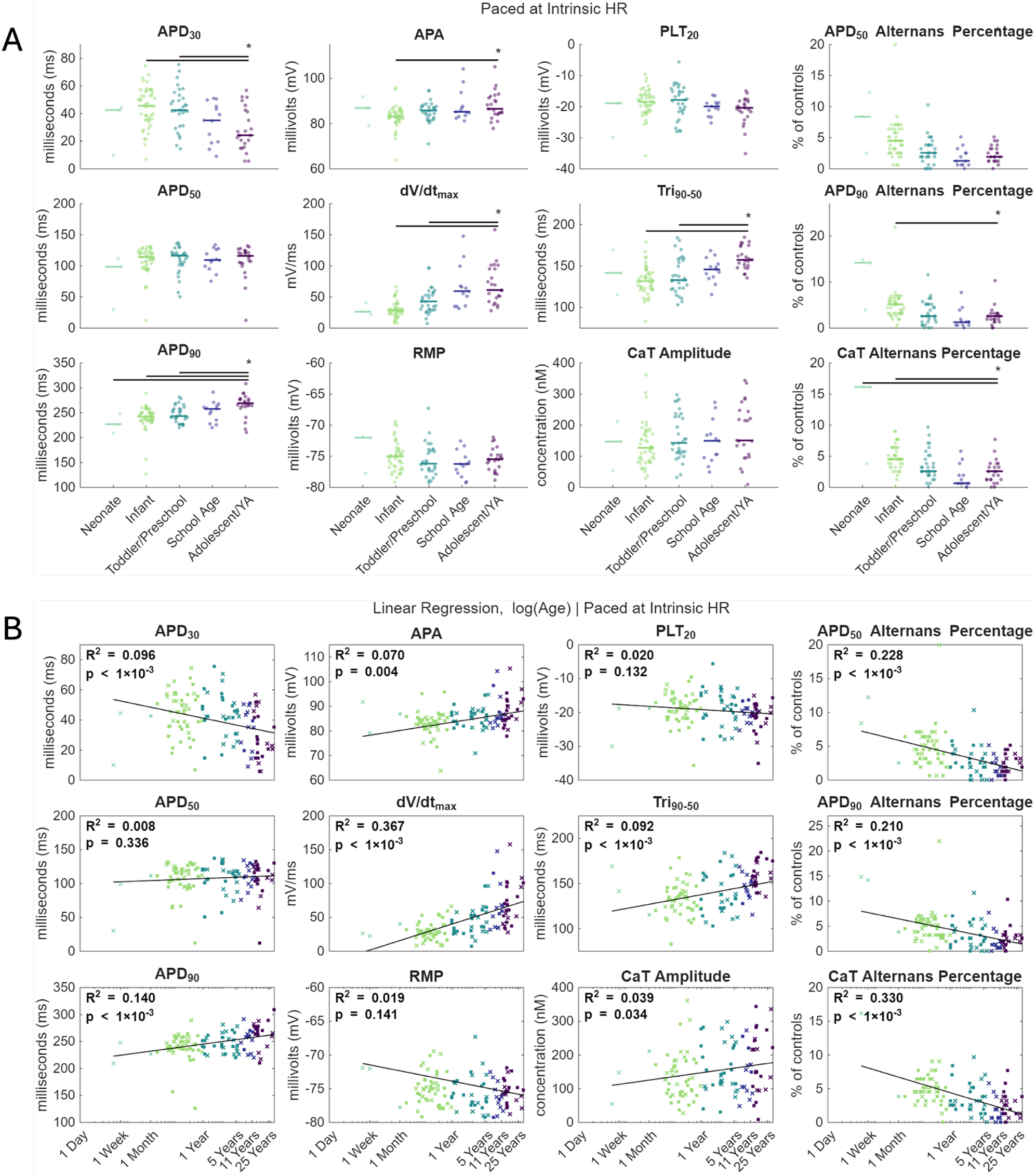
Age-dependent differences in simulated action potential and calcium transient biomarkers across pediatric development paced at intrinsic heart rate. A: Age group comparisons of action potential and calcium transient biomarkers presented as Beeswarm plots for each of the five age groups. Beeswarm plots also denote the median for each of the five age groups. Statistical comparison between each younger age group and the adolescent/young adult reference group were performed using one-way ANOVA when assumptions of normality and equal variance were satisfied, or Kruskal-Wallis tests otherwise, followed by Dunn-Šidák multiple comparisons; P < 0.05. B: Biomarkers plotted continuously as a function of age (days) on a logarithmic scale. Linear regression models were fit and the resulting regression lines, coefficient of determination (R^2^), and corresponding P values are shown. Each point represents an individual patient (males denoted by “□”, females by “×”), colored by age group. Abbreviations: APA, action potential amplitude; APD_30_, APD_50_, APD_90_, action potential duration at 30%, 50%, and 90% repolarization; CaT, calcium transient; dV/dt_max_, maximum depolarization rate; PLT_20_, plateau potential at 20% of APD_90_; RMP, resting membrane potential; Tri_90-50_, action potential triangulation.

### Influence of Pacing Rate on Developmental Electrophysiology

#### Comparison of Intrinsic and Fixed Pacing Conditions

To distinguish intrinsic developmental effects from rate-dependent influences, simulations were performed using each patient’s intrinsic heart rate and at fixed pacing rates of 64, 119, and 171 bpm, the patient cohort’s minimum, median, and maximum heart rates, respectively. Representative action potentials and calcium transients shown from one infant and one adult illustrate how the developmental stage and pacing rate influences simulated electrophysiological morphology (Figure 4, Supplemental Figure S1). For example, we see that through both the intrinsic and fixed pacing rates, the infant simulations tend to show shorter action potentials and larger amplitude CaTs, while the adult exhibits a more prolonged repolarization phase.

**Figure 4.**
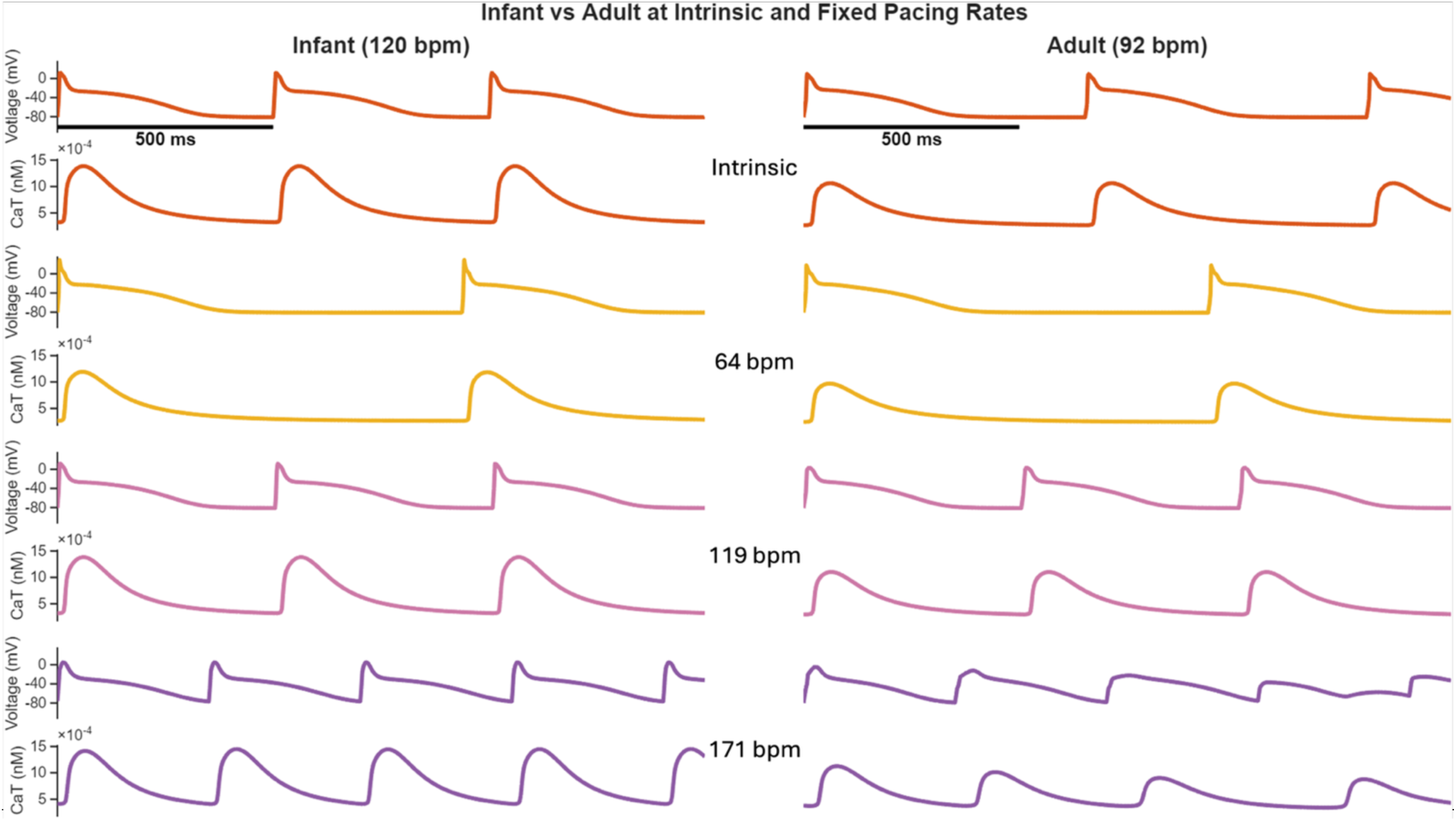
Simulated membrane voltage and intracellular calcium transients (CaT) are shown for a representative infant (left column; intrinsic rate of 120 bpm) and adult (right column; intrinsic rate of 92 bpm) cardiomyocyte models. For each age group, traces are presented at the intrinsic heart rate (top rows) and at fixed pacing rates of 64, 119, and 171 bpm (subsequent rows). Scale bars indicate 500 ms.

We next compared biomarker distributions within each age group across intrinsic and fixed pacing rates (Supplemental Figures S2-S9, Supplemental Table 2). Several rate dependent patterns were shared across age groups. Across all five age groups, faster pacing was associated with shorter repolarization-related biomarkers, including APD30, APD50, APD90, and Tri90-50, as well as slower dV/dt_max_, more elevated RMP, and increased prevalence in APD or CaT alternans (Supplemental Figures S2-S6). However, the magnitude of these pacing effects were not identical across developmental stages. Younger age groups generally exhibited a higher alternans burden than the adolescent/young adult group, whereas the older group showed lower alternans percentages across pacing conditions. Additionally, the distributions did not exhibit uniform increases in variability across all biomarkers. Greater spread was most apparent for alternans-related outcomes, including APD50 alternans percentage, APD90 alternans percentage, and calcium transient alternans percentage, whereas several electrophysiological biomarkers, such as APD30, APD50, APD90, dV/dt_max_, PLT20, RMP, and calcium transient amplitude, did not show consistently broader distributions in neonates or infants across pacing conditions. In general, for several electrophysiological biomarkers, pacing rate similarly affected simulated cellular response within each developmental group.

Further, several developmental trends were observed under both the intrinsic and fixed-rate pacing conditions (Supplemental Figures S7-S9). Across pacing conditions, earlier developmental stages exhibited shorter repolarization and greater alternans prevalence than the adolescent/young adult models, while later developmental stages exhibited greater dV/dt_max_ and more mature repolarization profiles. These trends indicate that age-dependent electrophysiological differences were not solely caused by the higher intrinsic heart rates of younger patients. However, the strength of age-dependent relationships differed between intrinsic and fixed pacing conditions. Under intrinsic heart rate pacing, age explained more variability (i.e., a larger R^2^) in dV/dt_max_ and alternans prevalence than under fixed pacing (Supplemental Table S2). In contrast, several APD metrics showed weaker or less consistent age associations across the fixed rate simulations. Together, these results suggest that developmental differences persist across pacing rates, but patient-specific intrinsic heart rate contributes substantially to age-associated differences in excitability and alternans prevalence.

#### Determinants of Electrophysiological Biomarkers

Finally, we used partial least squares (PLS) regression to quantify the relative contributions of ionic current conductances, calcium handling, and age to variability in simulated electrophysiological biomarkers (Figures 5-6). The PLS β-coefficient maps (Figure 5) demonstrated that each biomarker is influenced by a distinct combination of currents, however with both the magnitude and, in some cases direction, of these relationships varying across intrinsic and fixed pacing rates.

**Figure 5.**
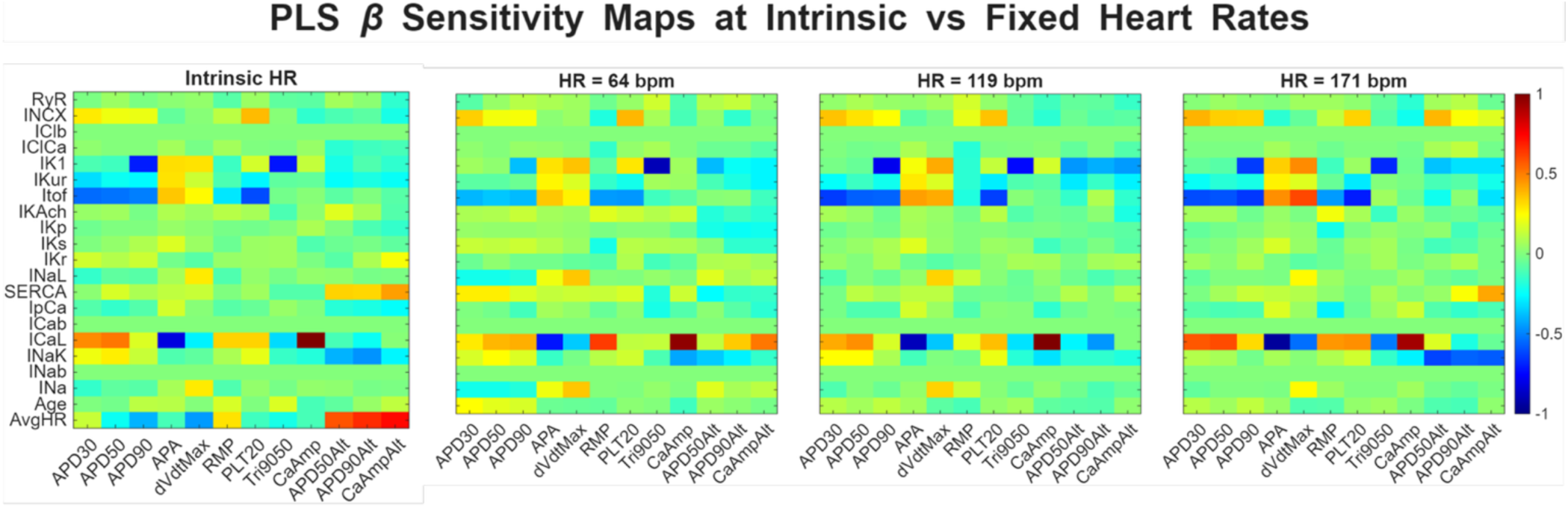
Partial least squares (PLS) regression β-coefficient sensitivity maps showing the influence of ionic current conductances and model parameters on simulated electrophysiological biomarkers under different pacing conditions. Subplots correspond to pacing conditions (intrinsic heart rate and fixed heart rates of 64, 119, and 171 bpm). Rows represent model parameters including ionic currents and calcium handling processes, while columns represent electrophysiological biomarkers derived from simulated action potentials and calcium transients. Colors indicate the magnitude and direction of the normalized β-coefficients, with warm colors representing positive associations and cool colors representing negative associations between parameters and biomarkers.

**Figure 6.**
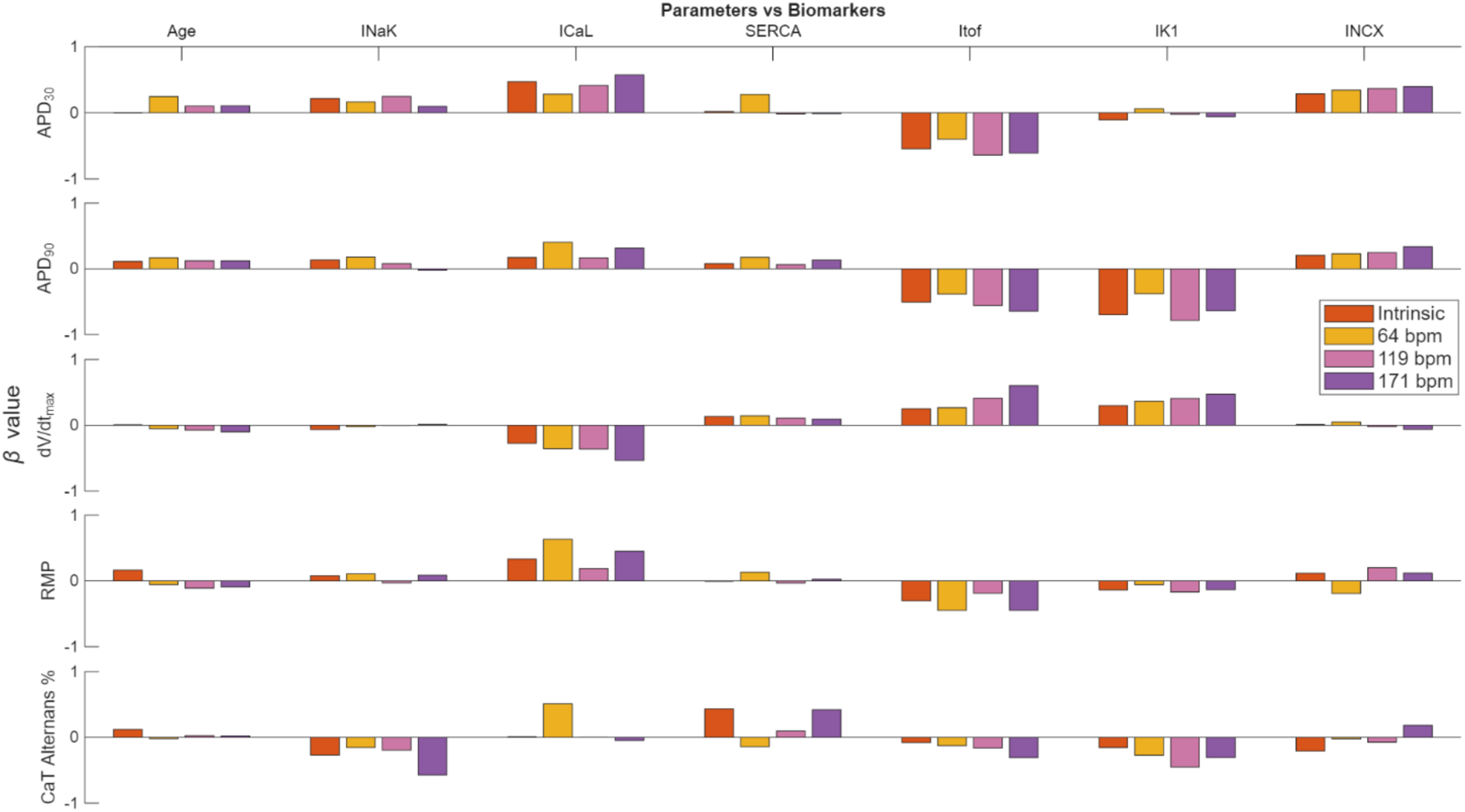
Partial least squares (PLS) regression β-coefficients illustrating the influence of selected model parameters on electrophysiological biomarkers across pacing conditions. Columns represent model parameters, including age and key ionic current conductances, while rows correspond to simulated biomarkers. Bars show the normalized β-coefficients obtained from PLS regression for intrinsic heart rate pacing and fixed pacing rates of 64, 119, and 171 bpm. Positive and negative β values indicate the direction and relative strength of each parameter’s contribution to the corresponding biomarker.

Repolarization-related biomarkers, including APD50, APD90, and action potential triangulation, were consistently associated with potassium currents, particularly IK1, IKur, and Itof. As expected, across heart rate conditions, IK1 exerted a strong influence on late repolarization and action potential termination (i.e., APD90), while IKur and Itof contributed more prominently to earlier phases of repolarization and shaping of the action potential plateau. Notably, the relative contribution of these currents shifted with pacing rate, with IK1 influence becoming more pronounced at faster pacing frequencies, consistent with an increased reliance on repolarization reserve under conditions of reduced diastolic recovery time. In contrast, early depolarization and upstroke velocity (dV/dt_max_) showed strong dependence on sodium current-related parameters, with additional modulation by background currents and electrogenic processes.

Plateau-phase characteristics, including PLT20 and intermediate repolarization dynamics, were strongly influenced by calcium and exchanger currents, particularly ICaL and INCX. ICaL contributed positively to plateau maintenance and calcium influx, while INCX demonstrated a prominent role in shaping plateau potential through its electrogenic exchange of calcium and sodium. The balance between these inward currents and opposing outward potassium currents was a key determinant of plateau morphology, with shifts in this balance across development contributing to observed age-dependent differences. CaT amplitude and duration were primarily governed by ICaL and SERCA activity, with SERCA contributing to calcium reuptake and influencing both the amplitude and kinetics of intracellular calcium cycling. Similarly, potassium currents exhibited differential contributions to repolarization metrics depending on pacing frequency, highlighting the rate-dependent redistribution of repolarization reserve.

Interestingly, across simulations paced at the intrinsic pacing rate, age is a fairly minimal independent determinant for most biomarkers, while average heart rate exhibits a stronger dependence. However, for fixed pacing rate conditions, age emerges as a moderate predictor, in particular for repolarization-related metrics, with the strongest dependence for APD30.

In Figure 6, we plot the normalized β-coefficients for subset of parameters (i.e., currents and age) and biomarkers to further illustrate the relative importance of specific parameters across biomarkers and pacing conditions. These results demonstrated that while certain currents consistently contributed to specific electrophysiological features, their relative weighting was not fixed and instead varied dynamically with pacing rate. As noted above, age emerged as a moderate independent contributor across multiple repolarization biomarkers. Thus, the persistence of age as a significant predictor indicates that developmental maturation is not fully explained by ionic currents or pacing rate changes alone.

## Discussion

While our prior transcriptomic-guided simulations of human atrial cardiomyocytes used a fixed pacing rate to avoid introducing frequency-dependent effects, it is well-established that heart rate changes markedly throughout pediatric development. In this study, we incorporated patient-specific intrinsic heart rate into simulations of pediatric atrial cardiomyocytes to determine how developmental maturation and pacing rate interact to shape predicted electrophysiological function. Intrinsic heart rate declined with age, with the highest rates in neonates and infants and progressively slower rates in older patients. Substantial interpatient variability within each age group, however, supported the use of patient-specific pacing rather than age group alone to represent physiological pacing conditions.

Gene expression-guided virtual populations predicted developmental changes in key simulated atrial electrophysiology biomarkers. APD50 and APD90 increased with age, whereas APD30 decreased, suggesting differential remodeling of early and late repolarization. In addition, dV/dt_max_ increased, resting membrane potential became more negative, and action potential triangulation increased with age. Alternans prevalence decreased across development, with younger groups exhibiting more APD50, APD90, and CaT alternans than the adolescent/young adult group. Together, these findings indicate coordinated postnatal changes in repolarization, excitability, and calcium handling.

Fixed-rate simulations showed that these developmental differences were not explained solely by the faster intrinsic heart rates of younger patients. Several age-dependent features, including shorter repolarization and greater alternans prevalence in younger groups, persisted across pacing conditions. Nevertheless, pacing rate modified their magnitude: faster pacing generally shortened repolarization-related biomarkers and increased alternans across age groups. Partial least squares regression further indicated that these outputs arose from distributed interactions among ionic conductances, calcium handling parameters, age, and heart rate rather than from a single dominant determinant. Pediatric atrial electrophysiology therefore appears to reflect both intrinsic developmental remodeling and rate-dependent modulation, with patient-specific heart rate contributing particularly to age-associated differences in excitability and alternans susceptibility.

The computational framework extends the transcriptomic-guided approach of our prior work, Salameh et al., where we incorporated pediatric right atrial gene expression data to a human atrial cardiomyocyte model to predict developmental differences in action potential and CaT morphology (24). Rather than using a single cell to represent each patient, we represented each patient with a virtual population generated by scaling conductances according to normalized gene expression relative to a baseline or control adult population. We retained this patient-specific population structure because direct electrophysiological recordings from pediatric human atrial myocytes are limited and a single model parameter set does not capture the observed range of patient-level variability.

This strategy is consistent with the broader shift in computational cardiac electrophysiology from single “average” cells toward the population-of-models approach that preserve variability in ionic current expression and electrophysiological phenotype. In adult human right atrial myocytes, Muszkiewicz et al. integrated experimental action potential and ionic current measurements with populations of human atrial models, demonstrating that experimentally calibrated populations reproduced action potential and calcium transient variability more effectively than a single baseline model (27). Population-based ventricular simulations have likewise been used to predict drug-induced proarrhythmic risk, identify vulnerable ionic profiles, and explain incomplete penetrance in inherited arrhythmia syndromes (28–31). More recently, virtual population and virtual patient approaches have been applied to atrial fibrillation, where interpatient differences in ionic currents, anatomy, and remodeling can influence predicted therapeutic responses (32,33). Collectively, these studies support population-based simulation as a means of preserving biological heterogeneity and identifying phenotypes that may be obscured by a single deterministic model.

Our approach differs from patient-specific strategies that estimate ionic conductances by fitting simulated action potentials directly to experimental recordings or through optimization and data-assimilation methods (34–36). Such methods can tightly constrain model behavior, including responses across pacing rates, when patient- or cell-specific electrophysiological recordings are available. Because these measurements are rarely obtainable from pediatric human atrial tissue, we instead used transcriptomic-guided conductance scaling to connect patient-level molecular data with predicted electrophysiological function. This strategy resembles prior efforts to inform personalized electrophysiological models with mRNA expression profiles (34,37). As mRNA levels do not necessarily correspond directly to protein abundance, channel trafficking, or current density, these simulations should be interpreted as mechanistic predictions rather than direct reconstructions of individual patients’ action potentials, which is an acknowledged limitation of our approach.

The principal methodological extension was the incorporation of patient-specific intrinsic heart rate. Salameh et al. paced all simulations at 1 Hz, or 60 beats/min, to isolate developmental effects derived from gene expression-based conductance scaling without introducing frequency dependence (24). Although appropriate for that objective, a fixed pacing rate does not establish how developmental variation in intrinsic heart rate influences electrophysiological phenotype. Simulating each virtual population at the corresponding patient’s intrinsic rate more closely approximated the physiological pacing environment, while simulations across fixed rates spanning the cohort range separated features that persisted across pacing conditions from those primarily modulated by rate.

This cellular-scale focus also distinguishes the study from broader pediatric cardiovascular models emphasizing multiscale digital twins, patient-specific anatomy, computational fluid dynamics, or clinical ECG-based parameter estimation (38,39). Those approaches address organ-level anatomy, flow, conduction, and clinical signals, whereas the present framework isolates the interaction between developmental gene expression-derived conductance differences and pacing rate at the single-cell level. It does not reconstruct whole-heart electrophysiology or tissue-level arrhythmia mechanisms, but it enables controlled assessment of how heart rate modifies action potential morphology, CaT behavior, and alternans susceptibility within patient-specific model populations.

The results support and refine the central conclusion of Salameh et al. that age is an important determinant of simulated pediatric atrial electrophysiology. Salameh et al. identified age-dependent differences in APD30, APD50, action potential triangulation, and plateau potential and reported strong associations of INCX with plateau potential and IK1 with triangulation (24). In the present study, differences between younger and adolescent/young adult models persisted under both intrinsic- and fixed-rate pacing, confirming that developmental electrophysiological variation is not solely a consequence of heart rate. Incorporating patient-specific rate, however, altered the magnitude and, for some biomarkers, the interpretation of these differences. Thus, the fixed-rate simulations of Salameh et al. isolated gene expression-dependent effects under a common pacing condition, whereas the current simulations show how those conductance differences are expressed within the physiological rate environment of pediatric patients.

This distinction was particularly evident in repolarization. Salameh et al. reported a developmental decrease in plateau potential and an increase in action potential triangulation, consistent with maturation-related changes in action potential morphology (24). We similarly observed developmental differences in repolarization across pacing conditions, but APD30, APD50, APD90, and triangulation also changed with rate. Under intrinsic-rate pacing, younger patient-specific models exhibited shorter repolarization and greater alternans than adolescent/young adult models. These findings complement rather than contradict the fixed-rate results: fixed-rate simulations isolate conductance-based maturation, whereas intrinsic-rate simulations demonstrate how that maturation is expressed under age-appropriate pacing.

The results also extend our previous calcium-handling findings that identified remodeling involving INCX as an important contributor to developmental differences in plateau potential (24). Our sensitivity analyses similarly implicated calcium and exchanger currents, including ICaL, INCX, and SERCA-related cycling, in plateau-phase behavior and calcium transient biomarkers. The intrinsic- and fixed-rate comparisons further showed that these outputs were rate sensitive, with faster pacing altering calcium transient amplitude and alternans, particularly in younger groups. Developmental remodeling of calcium handling therefore appears to establish the underlying phenotype, while heart rate modulates its functional expression.

The two studies also differed in their assessment of arrhythmia-related outputs. Salameh et al. evaluated abnormal action potentials and found that their simulated arrhythmia score had only modest predictive value for documented clinical arrhythmias, without a clear age-specific trend (24). We instead examined APD50, APD90, and CaT amplitude alternans as rate-sensitive indicators of electrophysiological instability. Their stronger age associations under intrinsic-than fixed-rate pacing suggest that patient-specific rate contributes to developmental differences in alternans susceptibility. Because electrical and calcium alternans are rate dependent and can create conditions favorable to arrhythmia, these outputs should be regarded as mechanistic markers of cellular instability rather than direct predictors of clinical arrhythmia (22,23).

More broadly, the developmental patterns support the view that the pediatric atrium is a distinct, rate-adapted electrophysiological state rather than simply an adult atrium paced more rapidly. The shorter APD50 and APD90, less negative resting membrane potential, and lower dV/dt_max_ in younger models are consistent with reported developmental differences in human atrial action potential morphology, transient outward potassium current, and L-type calcium current (12,15,18,20). Their persistence during fixed-rate pacing indicates that they arise at least partly from intrinsic molecular maturation. Faster pacing shortened repolarization across age groups, accommodating shorter cycle lengths, but also increased action potential and calcium transient alternans. The particularly high alternans burden in younger models may therefore reflect reduced stability of repolarization and calcium cycling during rapid activation, not simply their higher physiological heart rates.

These findings may have implications for age-appropriate pharmacology because responses to ion-channel-active drugs depend on the currents supporting repolarization, excitability, and calcium homeostasis and on the pacing rate at which those currents operate. Drugs that inhibit IKr, INa, or ICaL or alter intracellular calcium cycling may therefore produce different electrophysiological effects in infants and adolescents even after exposure is adjusted for body size. The present study identifies potential age- and rate-dependent pharmacodynamic differences that require further computational prediction and experimental testing. Recent pediatric studies of sotalol in early infancy and landiolol across a broad pediatric age range demonstrate the feasibility and value of evaluating efficacy, QT effects, safety, and pharmacokinetics directly in children (40,41). In patients receiving antiarrhythmic or other QT-active medications, developmental electrophysiology and intrinsic heart rate may therefore merit consideration alongside electrolyte abnormalities, concurrent medications, and perioperative stress when determining the need for ECG and rhythm monitoring.

Several limitations of the present study should be considered. First, as noted above, the framework assumes that normalized mRNA expression is proportional to ionic conductance and therefore does not directly represent protein abundance, channel trafficking, post-translational regulation, or functional current density. Second, the single-cell simulations omit cell–cell coupling, atrial anatomy, conduction heterogeneity, fibroblast interactions, dynamic autonomic regulation, and tissue-level reentry. Third, tissue was obtained from patients with acyanotic congenital heart disease, so the observed patterns may not fully represent healthy pediatric atrial maturation. Accordingly, the simulated action potentials, CaT, and alternans should be interpreted as mechanistic predictions rather than direct reconstructions of individual electrophysiology or clinical arrhythmia risk. These limitations motivate experimental validation, tissue-level modeling, and incorporation of age- and rate-dependent pharmacological responses.

As suggested above, future work with the present modeling approach can inform drug responsiveness and associated drug-induced risk. As such, a major direction for future work is to incorporate drug interactions into this patient-specific, heart-rate-dependent framework. As preliminary proof of concept simulations, β-adrenergic signaling was stimulated in representative infant- and adult-derived models, with each conductance profile simulated at an infant heart rate of 134 beats/min and an adult heart rate of 92 beats/min (Figure 7). Simulated isoproterenol altered action potential morphology and calcium cycling in all four combinations of developmental profile and pacing rate, including qualitative changes in plateau and repolarization trajectory, calcium transient amplitude, and diastolic calcium. The magnitude and time course of these responses differed between developmental profiles and pacing conditions, illustrating that pharmacological effects may depend jointly on molecular maturation and the heart rate at which the myocardium operates. Future studies will apply drug formulations across the complete patient-specific populations, quantify effects on repolarization, calcium handling, and alternans susceptibility, and investigate the extent to which particular developmental stages and/or ionic profiles are more vulnerable to drug-induced electrophysiological instability.

**Figure 7.**
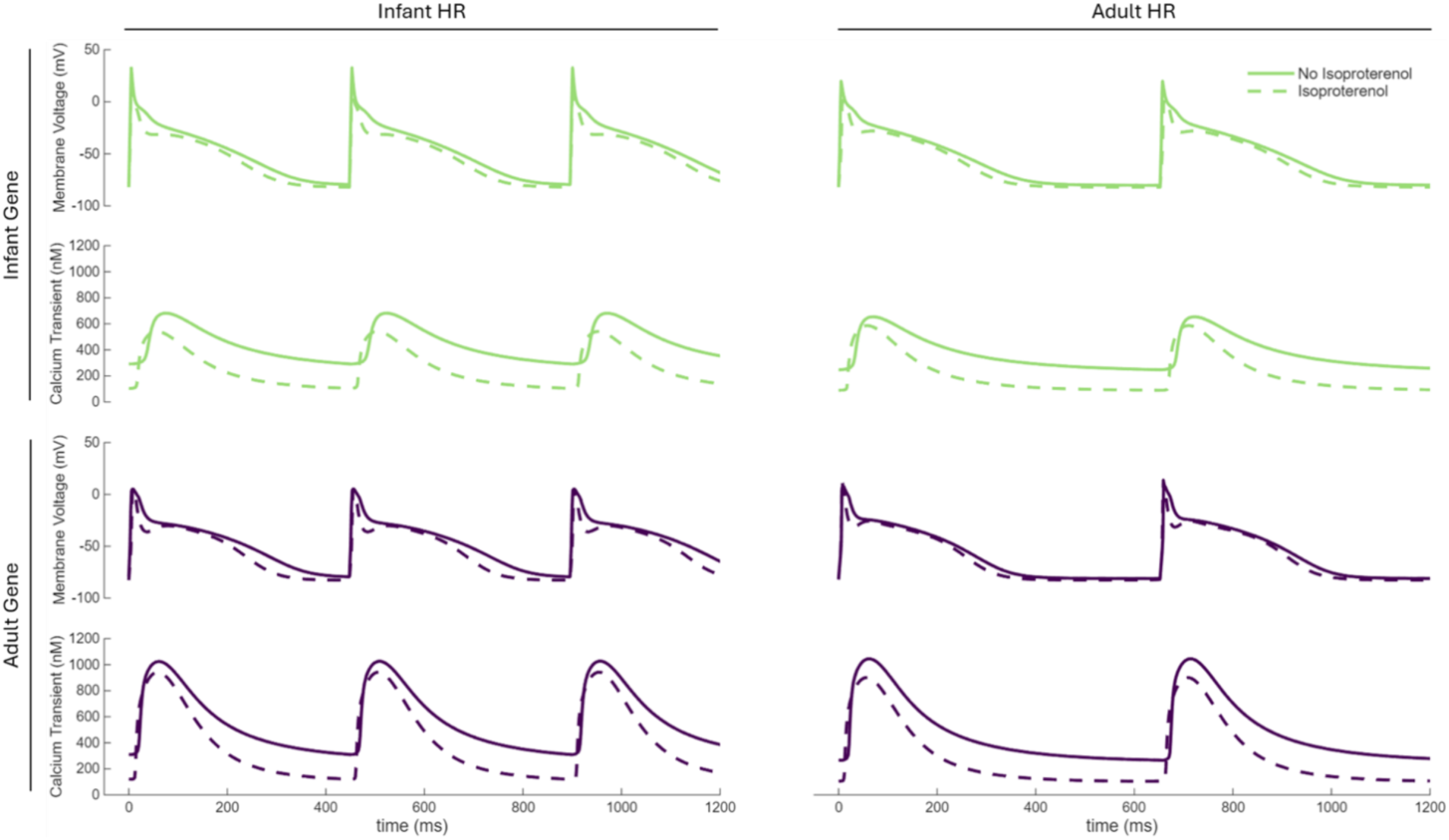
Proof-of-concept simulations of developmental and heart rate dependent responses to isoproterenol. Simulated membrane voltage and intracellular calcium transient traces are shown for a representative infant-derived (green, top) and adult-derived (purple, bottom) gene-expression based conductance profiles. Each profile was simulated at the associated representative infant-intrinsic heart rate of 134 beats per minute (left) and adult-intrinsic heart rate of 92 beats per minute (right). Solid traces represent control simulations without isoproterenol, and dashed traces represent simulations with the binary isoproterenol input activated using the model’s existing β-adrenergic signaling formulation.

## Acknowledgements

This work was supported by National Institutes of Health Grants R01HD108839 (to N.G.P. and S.H.W.), P30HD040677 (to C.N.R.I.), R01HL165751 (to S.H.W.), and R01HL169610 (to S.H.W.); Children’s National Heart & Lung Center, philanthropic support from the Seelig family, and the Foglia-Hills Professorship in Pediatric Cardiac Research.

We acknowledge additional members of the cardiac surgery operating room team, including Aybala Tongut, Alisa Bruce, Blezzy Bote, Alecia Byrd, Jannette Callos, Kenisha Cyrus, Moozhda Hanif, Evelyn Ravizee, Sandra Sunderland, and Hyung Mi (Grace) Yang for their assistance with tissue sample procurement. We also acknowledge Hanane Bousfoul and Riley Gardiner for their assistance with electrocardiogram waveform extraction.

