## Supplemental Tables and Figures for "Patient-Specific Heart Rate Modulates Developmental Electrophysiology in Transcriptomic-Guided In Silico Models of Pediatric Human Atrial Cardiomyocytes"

**SUPPLEMENTAL MATERIAL**

**Supplemental Tables**

**Supplemental Table S1.** *Patient Demographics*

| Age Group | Neonate | Infant | Toddler/Preschool | School Age | Adolescent/Young Adult |
|---|---|---|---|---|---|
| Number of Patients (Females) | 3 (2) | 47 (21) | 31 (14) | 13 (10) | 23 (9) |
| Age Min (days) | 5 | 41 | 366 | 2347 | 4396 |
| Age Max (days) | 30 | 361 | 2103 | 4373 | 11745 |
| HR Min (bpm) | 150 | 114 | 81 | 76 | 64 |
| HR Max (bpm) | 171 | 157 | 152 | 116 | 122 |

*Demographic and clinical characteristics of the study cohort stratified by pediatric age group, including number of patients, age range (days), heart rate (HR) range (beats per minute), and sex distribution.* Tissue samples were designated to one of five age groups: neonate (0–30 days, *n* = 3), infant (31–364 days; *n* = 47), toddler to preschool (1–5 yr, *n* = 31), school age (6–11 yr, *n* = 13), and adolescent to young adults (12–32 yr, *n* = 23). Note in this study, we used patient's chronological age, as opposed to the corrected gestational age used in our prior study (24).

**Supplemental Table S2.** *Coefficients of determination (R²) from linear regression analyses*

| **Biomarker** | **$R^2$** | | | |
|---|---|---|---|---|
| | | Fixed at | | |
| | Intrinsic HR | 64 bpm | 119 bpm | 171 bpm |
| $APD_{30}$ | 0.096 | 0.161 | 0.107 | 0.069 |
| $APD_{50}$ | 0.008 | 0.074 | 0.035 | 0.039 |
| $APD_{90}$ | 0.140 | 0.009 | 0.020 | 0.025 |
| APA | 0.070 | 0.055 | 0.030 | 0.031 |
| $dV/dt_{max}$ | 0.367 | 0.085 | 0.106 | 0.100 |
| RMP | 0.019 | 0.015 | 0.006 | 0.024 |
| $PLT_{20}$ | 0.020 | 0.025 | 0.054 | 0.089 |
| $Tri_{90-50}$ | 0.092 | 0.020 | 0.004 | 0.010 |
| CaT Amp | 0.039 | 0.047 | 0.034 | 0.027 |
| $APD_{50}$ Alternans Percentage | 0.228 | 0.015 | 0.030 | 0.011 |
| $APD_{90}$ Alternans Percentage | 0.210 | 0.003 | 0.003 | 0.001 |
| CaT Alternans Percentage | 0.330 | 0.008 | 0.035 | 0.015 |

*Coefficients of determination (R²) from linear regression analyses of action potential and calcium transient biomarkers as a function of age across different pacing conditions, including intrinsic heart rate and fixed pacing rates (64, 119, and 171 bpm).*

### Supplemental Figures

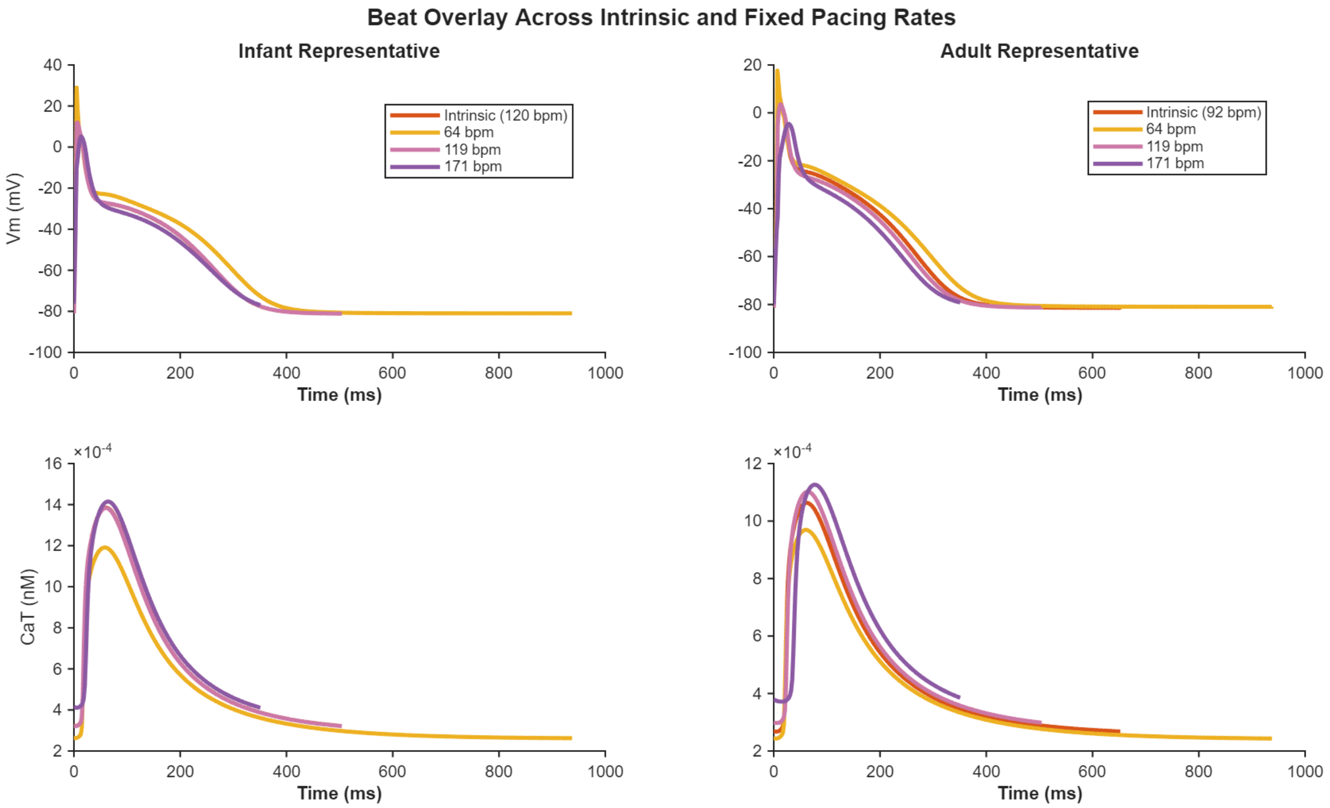


***Supplemental Figure S1.*** *Representative first-beat membrane voltage (Vm) and intracellular calcium transients (CaT) are overlaid for infant (left column; intrinsic rate of 120 bpm) and adult (right column; intrinsic rate of 92 bpm) cardiomyocyte models. Traces are shown for intrinsic pacing and fixed pacing rates of 64, 119, and 171 bpm, aligned to the onset of stimulation and plotted over a single cycle length for each condition.*

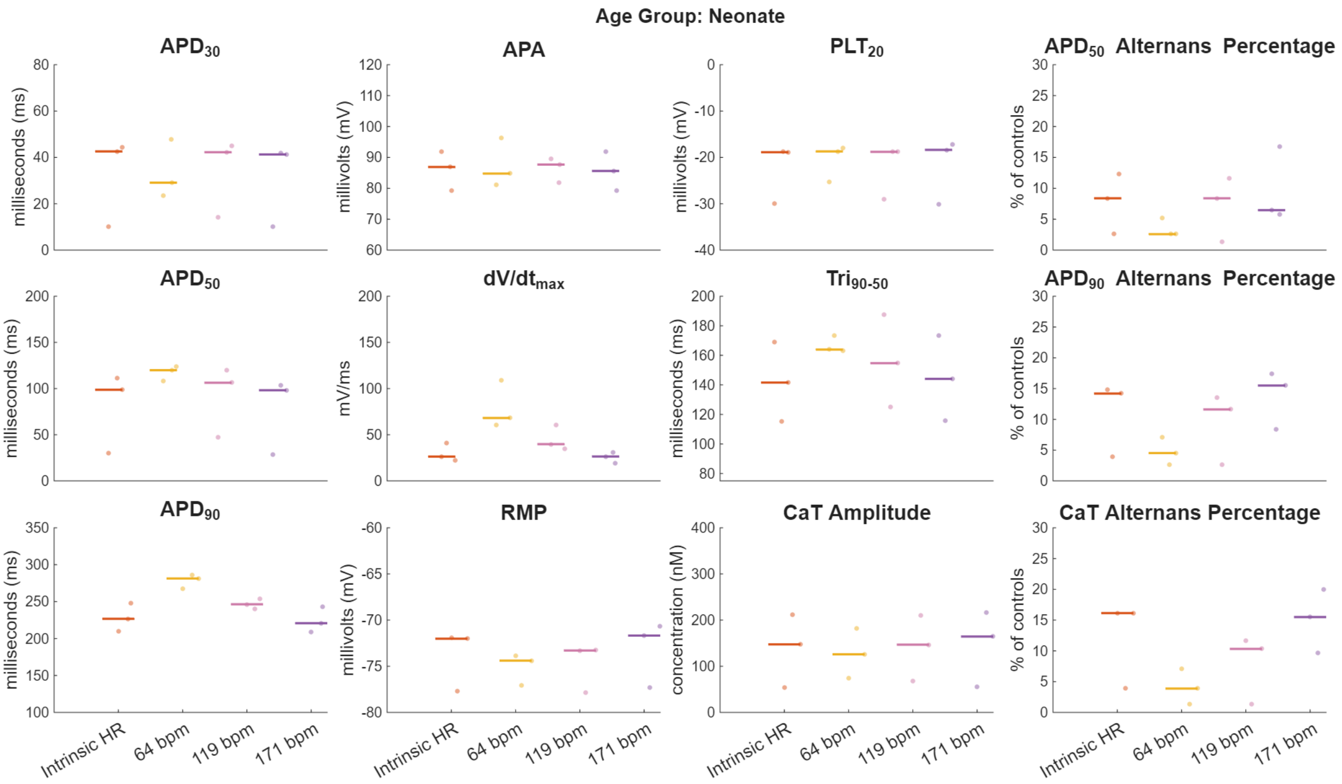


***Supplemental Figure S2.*** *Comparison of simulated action potential and calcium transient biomarkers in the neonatal age group across pacing conditions. Beeswarm plots show biomarker distributions under intrinsic heart rate pacing and fixed pacing rates of 64, 119, and 171 bpm.*

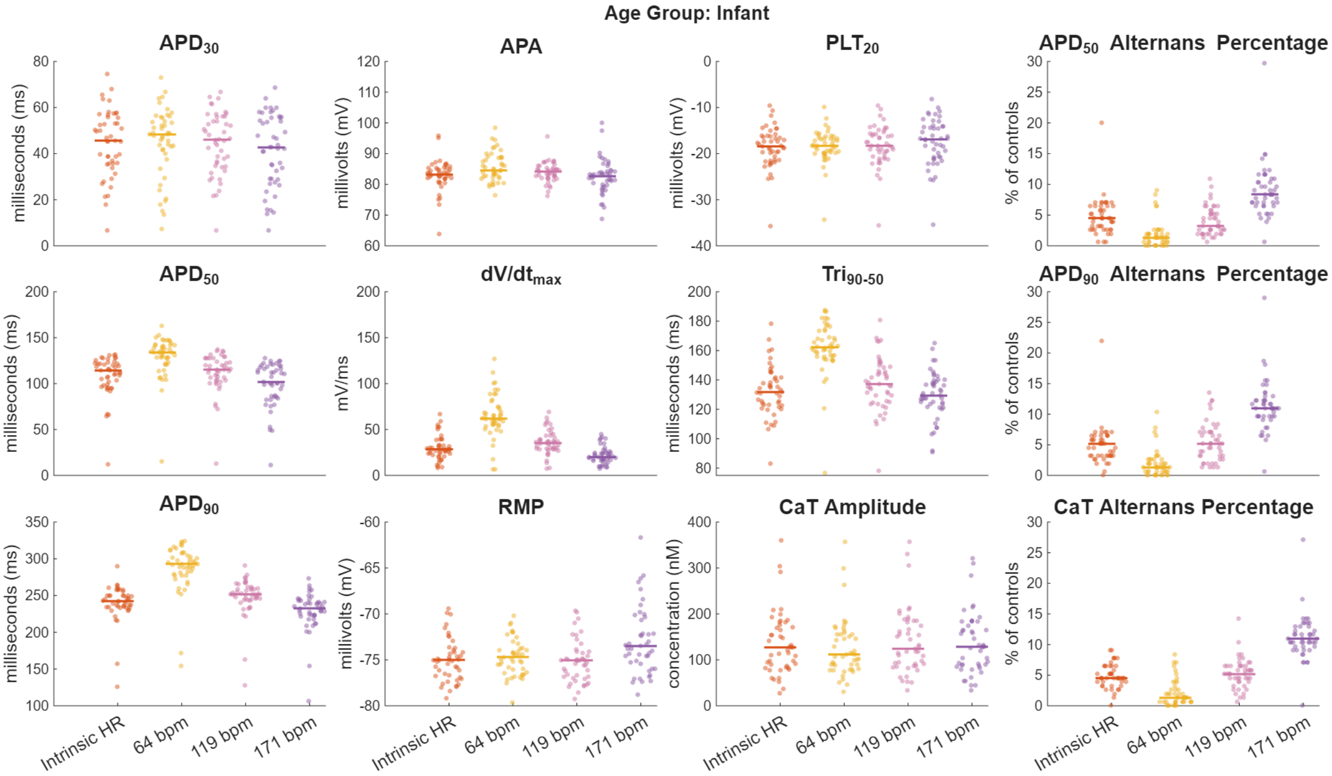


***Supplemental Figure S3.*** *Comparison of simulated action potential and calcium transient biomarkers in the infant age group across pacing conditions. Beeswarm plots show biomarker distributions under intrinsic heart rate pacing and fixed pacing rates of 64, 119, and 171 bpm.*

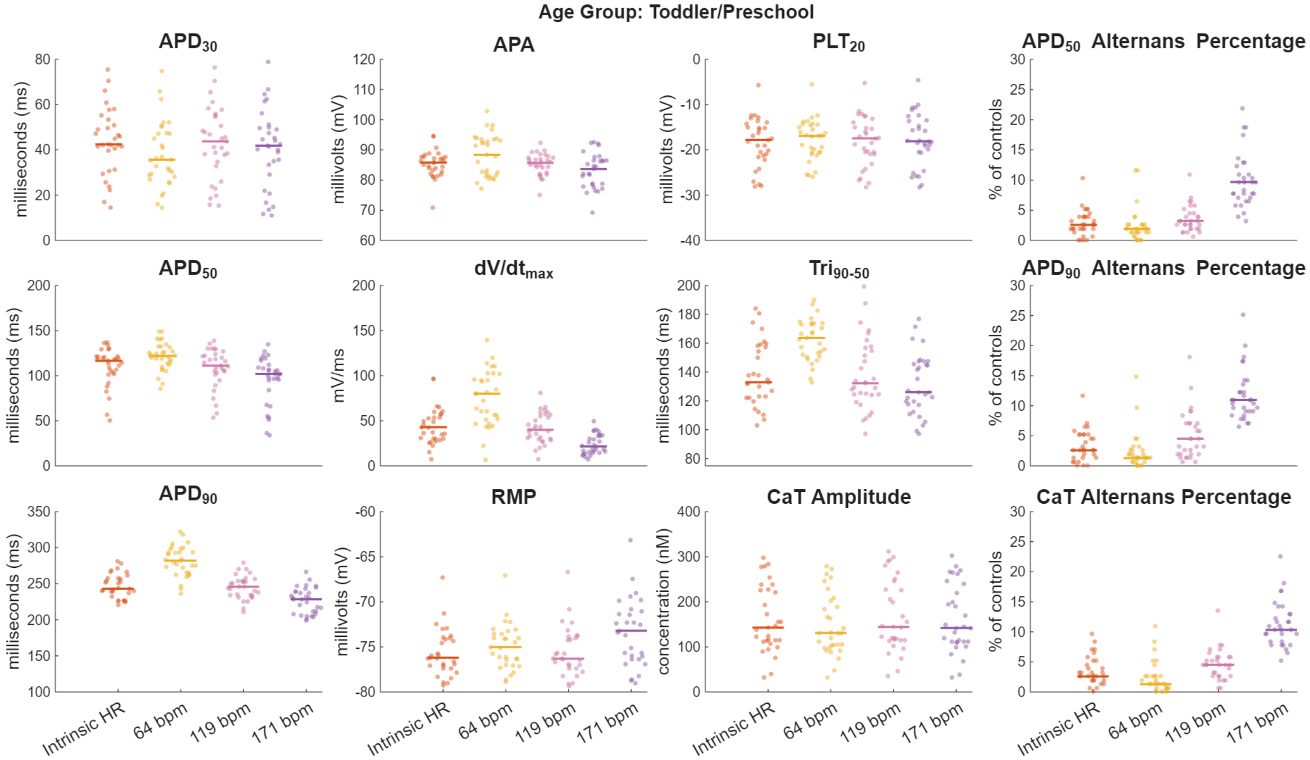


***Supplemental Figure S4.*** *Comparison of simulated action potential and calcium transient biomarkers in the toddler/preschool age group across pacing conditions. Beeswarm plots show biomarker distributions under intrinsic heart rate pacing and fixed pacing rates of 64, 119, and 171 bpm.*

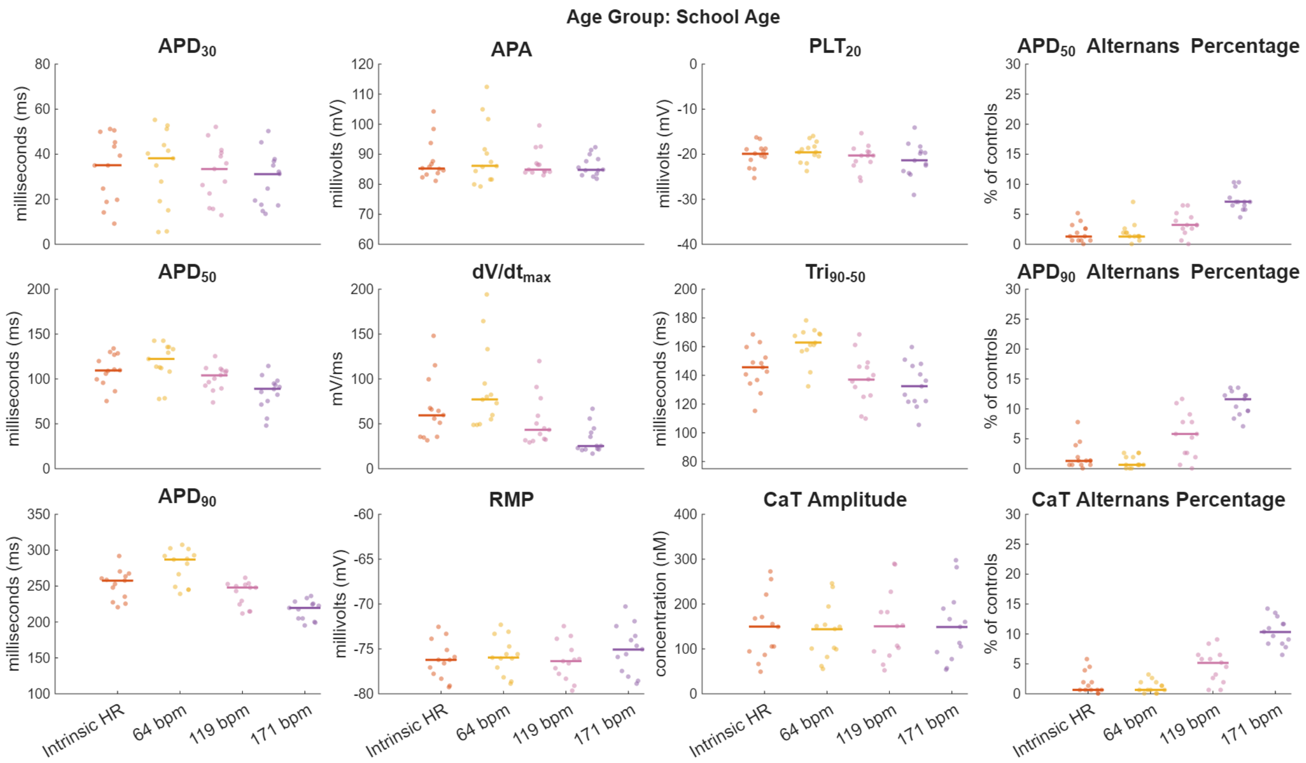


***Supplemental Figure S5.*** *Comparison of simulated action potential and calcium transient biomarkers in the school age group across pacing conditions. Beeswarm plots show biomarker distributions under intrinsic heart rate pacing and fixed pacing rates of 64, 119, and 171 bpm.*

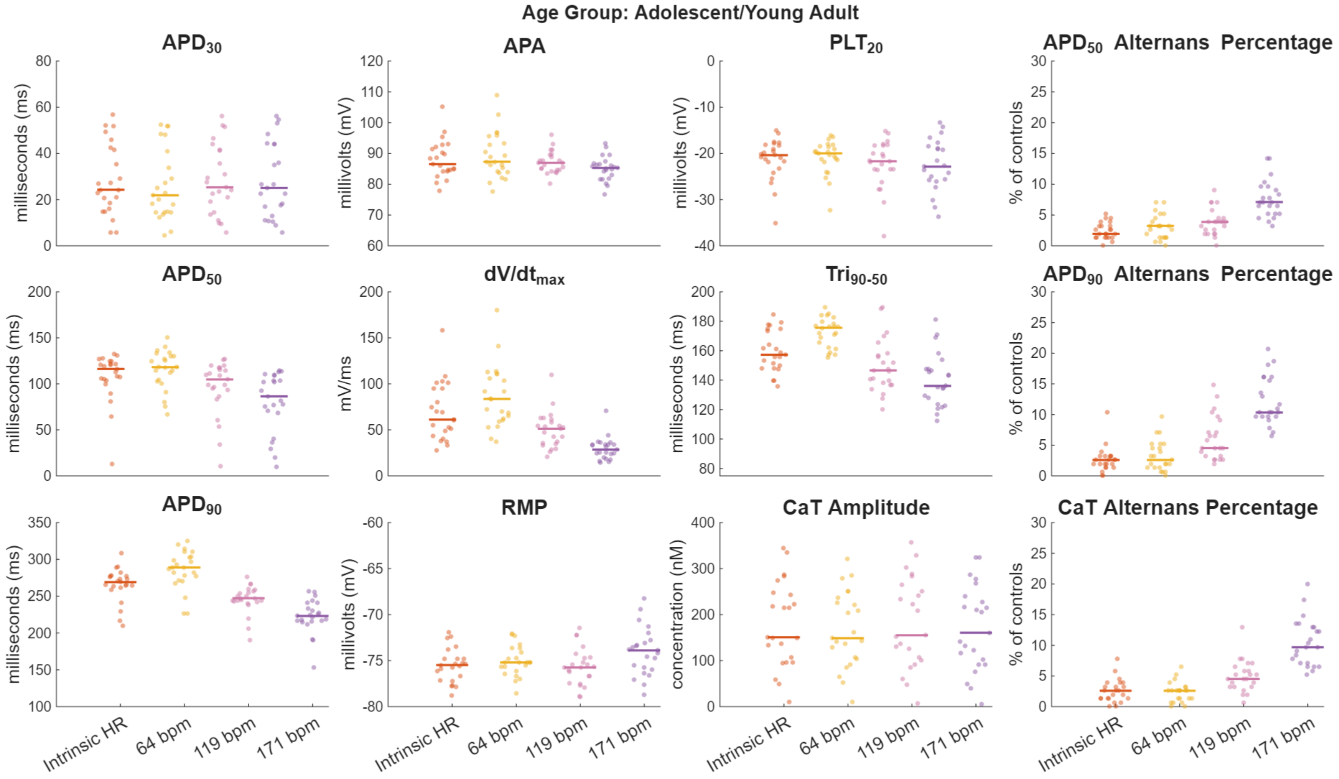


***Supplemental Figure S6.*** *Comparison of simulated action potential and calcium transient biomarkers in the adolescent/young adult age group across pacing conditions. Beeswarm plots show biomarker distributions under intrinsic heart rate pacing and fixed pacing rates of 64, 119, and 171 bpm.*

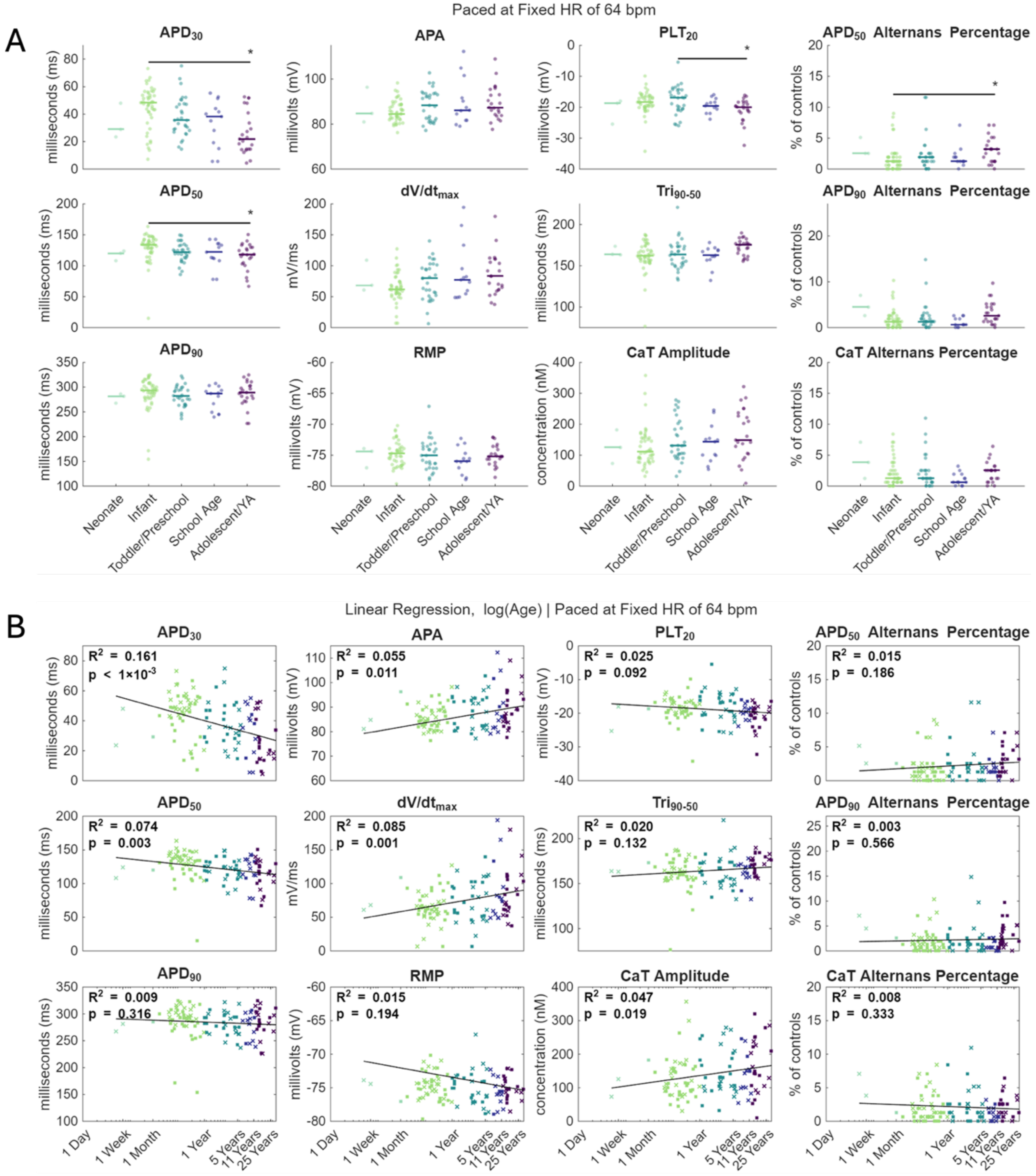


***Supplemental Figure S7.*** *Age-dependent differences in simulated action potential and calcium transient biomarkers across pediatric development paced at population minimum heart rate. Panel organization, statistical analysis, and regression methods are identical to those described in* ***Figure 3****, but simulations were performed at a constant heart rate of 64 beats per minute rather than the intrinsic heart rate of each age group.*

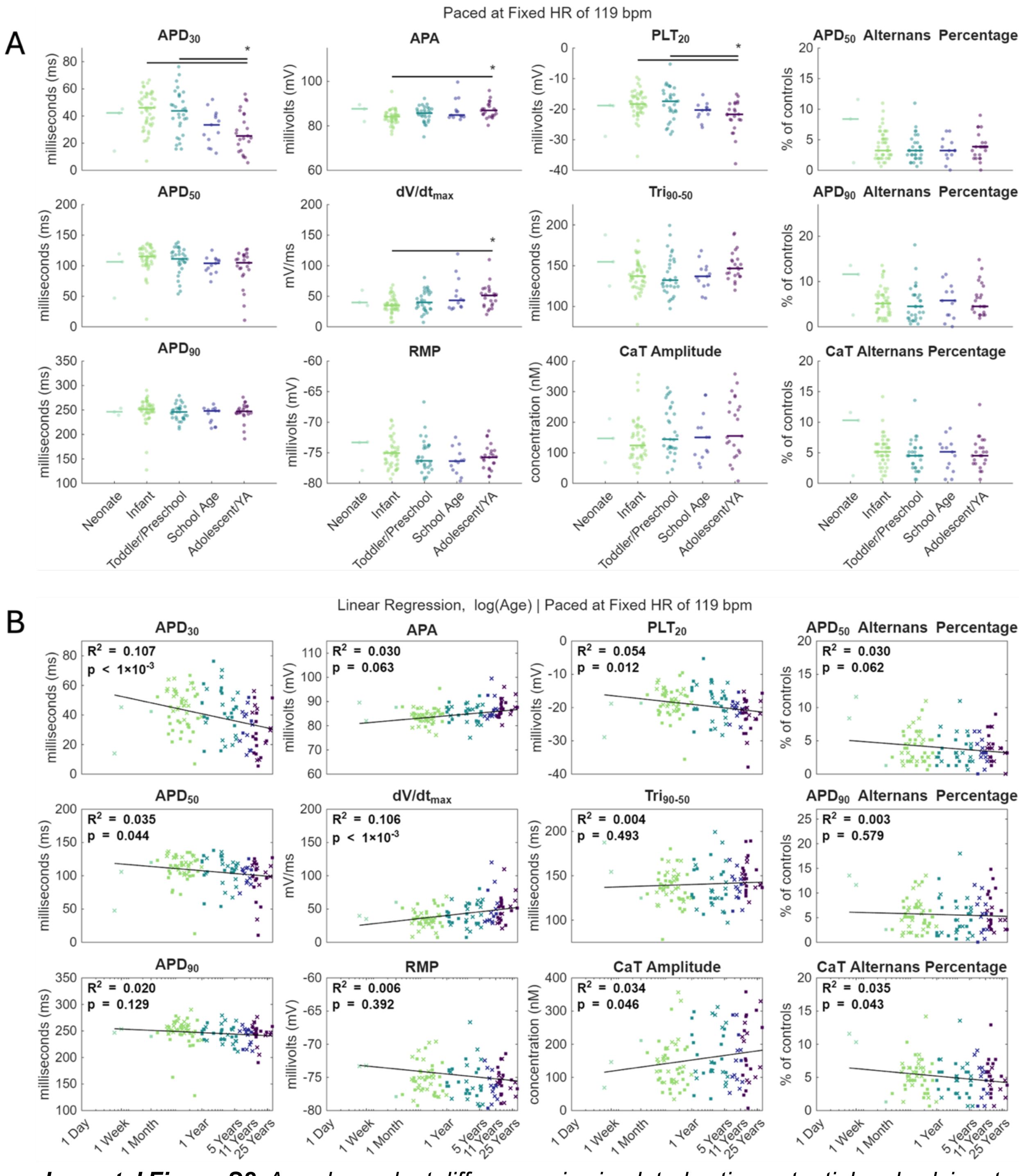


***Supplemental Figure S8.*** *Age-dependent differences in simulated action potential and calcium transient biomarkers across pediatric development paced at population median heart rate. Panel organization, statistical analysis, and regression methods are identical to those described in* ***Figure 3****, but simulations were performed at a constant heart rate of 119 beats per minute rather than the intrinsic heart rate of each age group.*

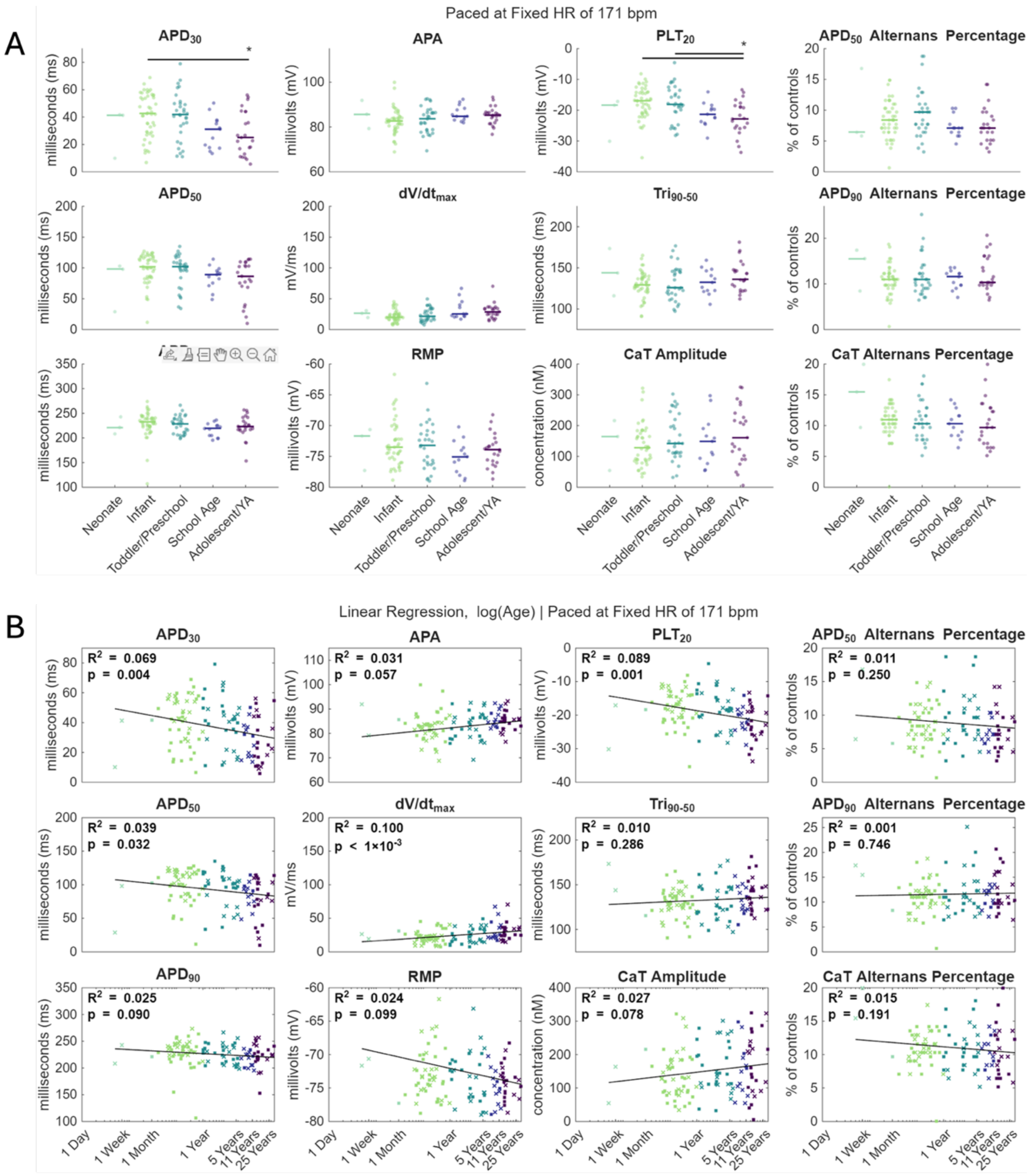


***Supplemental Figure S9.*** *Age-dependent differences in simulated action potential and calcium transient biomarkers across pediatric development paced at population maximum heart rate. Panel organization, statistical analysis, and regression methods are identical to those described in* ***Figure 3****, but simulations were performed at a constant heart rate of 171 beats per minute rather than the intrinsic heart rate of each age group.*
